# Correlative live and super-resolution imaging reveals the dynamic structure of replication domains

**DOI:** 10.1101/189373

**Authors:** Wanqing Xiang, M. Julia Roberti, Jean-Karim Hériché, Sebastian Huet, Stephanie Alexander, Jan Ellenberg

## Abstract

Chromosome organization in higher eukaryotes controls gene expression, DNA replication, and DNA repair. Genome mapping has revealed the functional units of chromatin at the sub-megabase scale as self-interacting regions called topologically associating domains (TADs) and showed they correspond to replication domains (RDs). A quantitative structural and dynamic description of RD behavior in the nucleus is however missing, as visualization of dynamic subdiffraction-sized RDs remains challenging. Using fluorescence labeling of RDs combined with correlative live and super-resolution microscopy *in situ*, we determined biophysical parameters to characterize the internal organization, spacing and mechanical coupling of RDs. We found that RDs are typically 150 nm in size and contain four co-replicating regions spaced 60 nm apart. Spatially neighboring RDs are spaced 300 nm apart and connected by highly flexible linker regions that couple their motion only below 550 nm. Our pipeline allows a robust quantitative characterization of chromosome structure *in situ*, and provides important biophysical parameters to understand general principles of chromatin organization.

## Introduction

The spatial arrangement of chromatin plays a fundamental role in genome function and stability (Dixon et al., 2016; Sexton et al., 2012; Gorkin et al., 2014; Phillips-Cremins et al., 2013). DNA replication and repair, gene transcription and cell differentiation depend on intra and inter-chromosomal contacts between genomic loci, promoter-enhancer interactions and accessibility of the DNA sequence to proteins (Gibcus et al., 2013). Specific chromosome regions that interact with nuclear structural elements, such as the nuclear envelope and nucleoli, provide an additional large scale structural layer to genome regulation and chromatin compartmentalization (Kind et al., 2013; Inoue and Zhang, 2014). To pack the genome in a mammalian cell nucleus of less than 10 μm diameter on average, chromosomal DNA molecules undergo multiple levels of compaction. The first occurs at the bp to Kbp scale via DNA-core histone association to nm-sized nucleosomes (Richmond et al., 1997). The last level is at the 100 Mbp scale, where whole chromosomes occupy defined µm-sized volumes inside the nucleus, termed chromosome territories (CTs) (Gilbert et al., 2004; Cremer and Cremer, 2001). The structures at the intermediate scale ranging from kbp to Mbp are not directly known *in situ*, but it is clear that subchromosomal chromatin domains with distinct epigenetic marks exist that are important to modulate gene expression (Bernstein et al., 2007). Over the last years, chemical cross linking-based genome-wide mapping methods, particularly HiC, have captured with unprecedented detail the frequency of contacts between linearly distant genomic loci and identified so-called topologically associating domains (TADs) as stable structural units of genome organization (Lieberman-Aiden et al., 2009). TADs have genomic sizes between 400 kbp and 1 Mbp (Dixon et al., 2012; Wang et al., 2016) and are separated by boundaries of ~50 kbp segments enriched in CTCF sites (Phillips and Corces, 2009), transfer RNA genes, and short interspersed nuclear elements serving as insulators for transcriptional regulation (Dixon et al., 2016). In size and number, TADs have many similar properties to the long known replication domains (RDs), which are co-replicating DNA sequences in the genome, that show very reproducible spacing and timing in many cell types and species (Rivera-Mulia and Gilbert, 2016; Jackson and Pombo, 1998). Very interestingly, the boundaries that separate TADs as mapped by HiC show an almost one-to-one correlation to the boundaries separating RDs as measured by replication analysis (Pope et al., 2014; Moindrot et al., 2012). This observation confirmed the long-standing notion that TADs in fact share replication timing and that TADs/RDs are the major organizational sub-chromosomal elements in eukaryotes.

While the genomic size of TADs/RDs and their spacing along the linear sequence of chromosomes have been well-defined, their physical size and internal structure has yet to be elucidated. Given the functional importance of these domains (Letourneau et al., 2014), complementary efforts are currently devoted to this task. Genome-wide biochemical mapping techniques typically infer higher-order chromatin interactions from averaging over cell populations, are not quantitative and lack direct spatial and temporal information. By contrast, fluorescence imaging can report direct physical spatial and temporal parameters. Especially fluorescence *in situ* hybridization (FISH) of specific genomic sequences has been very powerful, starting with the identification of CTs (Cremer et al., 1993) and the positioning of individual genes (Croft et al., 1999), and more recently the mapping of the spatial arrangements of TADs in human diploid cells by super-resolution microscopy (Wang et al., 2016).

Despite this considerable progress, many of the physical parameters to understand chromosome structure *in situ* are lacking. A particular challenge is the cross scales nature of the problem, necessitating not only to resolve the size and internal organization of TADs/RDs but also their physical distances as well as their dynamic relationships to each other within CTs. To address this gap in our knowledge, we have combined fluorescent labelling of single or neighboring RDs of one CT in living mammalian cells and quantitatively characterize them by correlative confocal and super-resolution microscopy *in situ*. This allowed us to address the internal organization of co-replicating regions inside RDs and estimate the physical size of these domains. In living cells, we could furthermore determine the spacing of adjacent RDs and reveal that the connection between them is highly flexible. Our data supports a model of chromosome organization, where 150 nm sized RDs with typically four co-replicating regions are spaced 300 nm apart on the chromosomal molecule and are linked by highly flexible linkers that couple their motion only below 550 nm. Our method is sequence independent and generic and allows a quantitative characterization of chromosome structure *in situ*. It provides important biophysical parameters missing to determine the organizational principles of chromosomes inside the nucleus of the cell.

## Results

### Fluorescence labeling of RDs for correlative confocal and super-resolution microscopy

We labeled the DNA backbone of euchromatic RDs using co-replicative pulse labeling with fluorescent nucleotides (Schermelleh et al., 2001), optimized for super-resolution microscopy by using appropriate fluorophores and gentle permeabilization, which resulted in a high survival rate and very good labelling efficiency of short DNA stretches that were replicating at the time of labeling (Fig. 1; for details see Materials and methods). After screening commercially available hydrophilic fluorophore-coupled nucleotides (Table S1), we chose ATTO 633-dUTP as the best dye for single-color confocal and correlative STORM imaging (Fig. 1B), and a 1:1 molar ratio combination of ATTO 565-dUTP and ATTO 633-dUTP as the best pair for dual-color live confocal microscopy (Fig. 1C). The use of this labeling protocol followed by several rounds of cell division resulted in cells containing only a few or single CTs with on average 22 fluorescently labeled RDs/CT (Fig. 1A, Fig. S1A-B), and a virtually background-free labeling (Fig. 1).

**Fig. 1.**
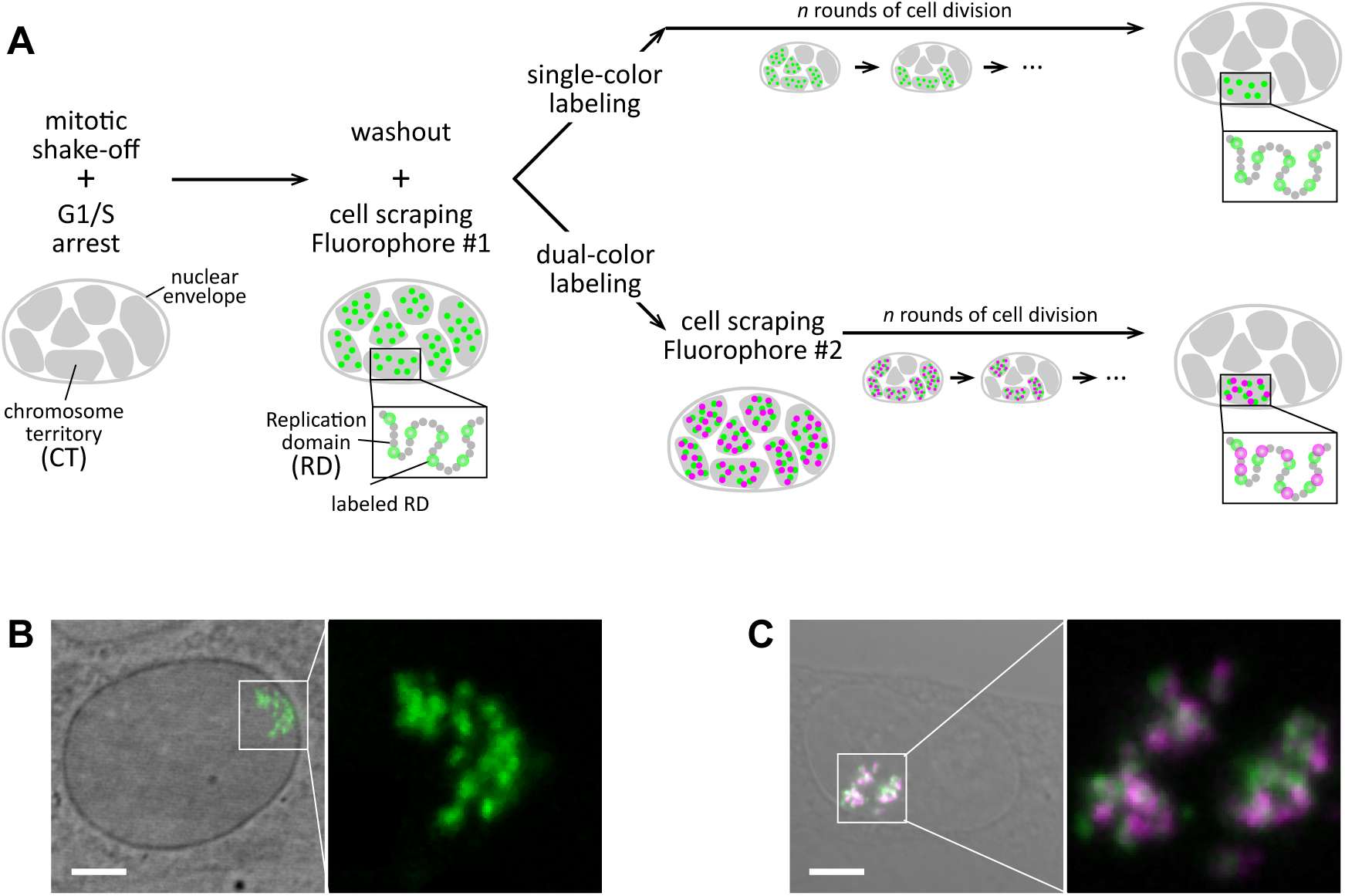
Co-replicative labeling of early-replicating RDs for correlative confocal-STORM imaging. (**A**) Schematics of the experimental approach. NRK cells were harvested from a mitotic shake-off and plated in the presence of aphidicolin to arrest them at the G1/S transition. After 10 h, aphidicolin was removed and replaced by culture medium containing the fluorophore of interest (Fluorophore #1, in this work ATTO633) coupled to dUTP. Cells were scraped off the culture chamber bottom, allowing the labeled dUTP to enter and be incorporated only in the actively co-replicating RDs (green circles), the rest remained unlabeled (gray circles). After labelling, cells underwent several rounds of division to allow a sparse localization of CTs within the nucleus (typically 3-4 days after the co-replicative labeling procedure). For dual-color experiments (bottom row), cells were subsequently scraped in the presence of a second fluorophore (Fluorophore #2, in this work ATTO565) coupled with dUTP to label actively co-replicating RDs at a different timepoint (green and magenta circles). The interval Δt separating the application of Fluorophore#1 and #2 was varied between 0 and 120 min. (**B, C**) Examples of cells with an ATTO633-dUTP single-color labelled CT (**B**) and with ATTO633- and ATTO565-dUTP double-color labeled CTs (**C**). Left panels: image of the nucleus with transmitted light (gray) overlaid with the fluorescence channel of labeled RDs (ATTO633 green, ATTO565 magenta); scale bar 5 µm. Right panels: 4x zoom-in of the fluorescence channel.

DNA combing (Shaw et al., 2010; Michalet et al., 1997) of ATTO633-dUTP labeled cells (Fig. S1C) revealed short, comet-like stretches of labeled DNA whose decreasing signal intensity reflected the consumption of the ATTO633-dUTP pool during progression of replication. Using stretched λDNA as a ruler, we estimated that we typically labeled co-replicating DNA stretches of 18.2 ± 4.5 kbp (Fig. S1D-F).

### Super-resolution microscopy reveals the internal structure of RDs which typically contain four co-replicating stretches

Using diffraction-limited confocal imaging with a lateral resolution of ~250 nm and an axial resolution ~750 nm, RDs appear as subdiffractive objects without discernable internal structure, and consequently previous estimates of RD size range from 350 nm (Shaw et al., 2010) to 500 nm (Albiez et al., 2006). However, indirect evidence suggested that RDs consist of small groups of replicons (Jackson and Pombo, 1998) which start DNA replication synchronously. Labeling with a pulse of fluorescent nucleotides therefore labels all actively replicating sequences within a RD. To reveal the organization of these co-replicating stretches inside RDs, we established single-cell correlative confocal and super-resolution imaging using stochastic optical reconstruction microscopy (STORM) of fluorescently labeled RDs (Fig. 2A, B). This allowed us to resolve the diffraction limited foci of single RDs into groups of discrete subdiffraction peaks (Fig. 2B,C) with a resolution better than 20 / 50 nm (estimated by Full width at half maximum (FWHM) and Fourier Ring correlation, respectively) and determine their positions by computational image analysis (for details see Material and methods). To characterize the number of co-replicating stretches per RD we applied the unbiased clustering algorithm DBSCAN (Density-based Spatial Clustering of Applications with Noise (Ester et al., 1996)), to the peak positions (Fig. 2D). Analysis of 9766 peaks and correlation to the confocal images revealed that 87% clustered into RDs in groups of two or more peaks while only 12.6% constituted presumably solitary co-replicating stretches that were undetectable as RDs by confocal imaging (Fig. 2D, yellow boxes). The distribution of the number of co-replicating stretches per RD followed an exponential decay, with a median number of four co-replicating stretches per RD (Fig. 2E).

**Fig. 2.**
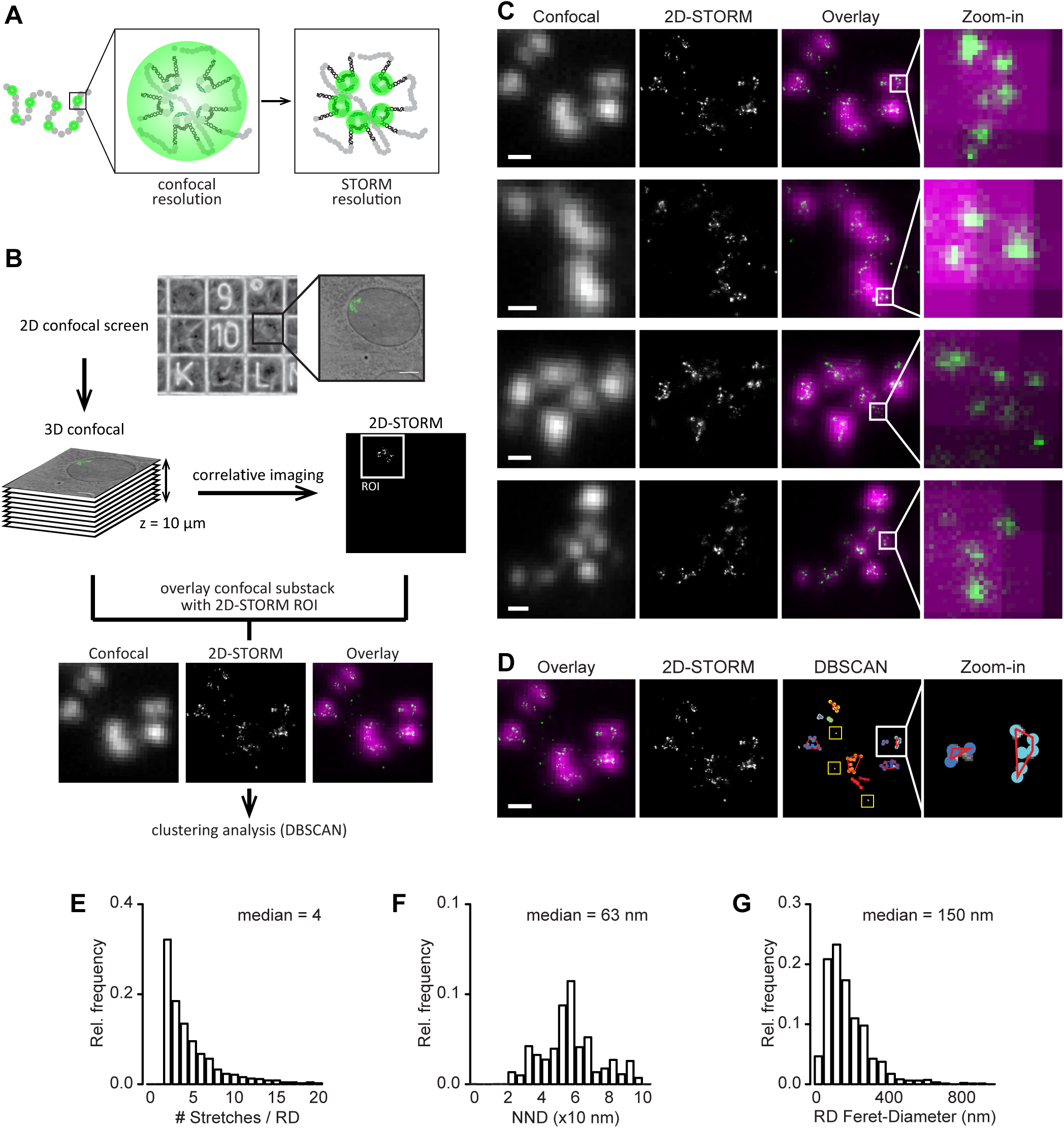
Correlative confocal-STORM imaging reveals the internal structure and size of RDs. (**A**) Cartoon of a single-color labeled CT. A subset of RDs is tagged by co-replicative labeling (green circles). Each RD consists of several replicons which are simultaneously activated during S-phase and co-replicate their DNA synchronously. At confocal resolution (ca. 300 nm), RDs appear as subdiffraction-sized spots, whereas at a resolution of ~20 nm using STORM imaging, it is possible to resolve the individual replicons or co-replicating stretches. (**B**) Experimental pipeline for correlative confocal-STORM imaging. NRK cells labeled with ATTO633-dUTP were plated onto gridded coverslips, screened in 2D to identify cells of interest containing a few CTs and to be imaged in 3D at confocal resolution. The cover slips were then put in switching buffer, and the cells of interest imaged in 2D with the STORM microscope at super-resolution. The correlative confocal stacks and 2D-STORM images were analyzed to find the optimal confocal substack-STORM overlay and to discard out-of-focus regions of the STORM images. Finally, DBSCAN clustering analysis was performed on the cropped STORM images to obtain estimates of number of co-replicating stretches/RD, RD size and RD diameter. (**C**) ATTO633-labeled CTs imaged by correlative confocal-STORM imaging. Panels from left to right: optimal confocal z-substack projection (gray), STORM image (gray), overlay (confocal in magenta, STORM in green), and 10x zoom-in. Scale bars: 500 nm. Small groups of co-replicating stretches can be resolved in the STORM images, and are not discernible from the confocal images. Number of CTs imaged *n* = 87, from 37 cells. (**D**) Representative Confocal-STORM overlay showing qualitatively the correspondence between diffraction-limited RDs and superresolved groups of co-replicating stretches. Clustering analysis: example STORM image after filtering and intensity thresholding; density-based clustering of detected co-replicating stretches to RD (color-coding shows forks belonging to the same cluster, and the red lines denote the convex hull around the centers of detected co-replicating stretches; yellow squares mark unclustered, solitary co-replicating stretches); scale bar 500 nm, and 5x zoom-in. (**E-G**) DBSCAN: histograms of the number of co-replicating stretches counted per cluster/RD and median value (**E**), the nearest neighbor distance (NND) between co-replicating stretches and median value (**F**), and the horizontal Feret-diameter of RD and median value (median including clusters of 2 co-replicating stretches = 105 nm) (**G**).

### RDs have a diameter of 150 nm with co-replicating DNA stretches spaced 63 nm apart

To characterize the spatial relationship of co-replicating stretches within a RD, we measured the nearest neighbor distances (NND) between STORM peaks and found a median distance of 63 nm (Fig. 2F). The cluster-based assignment of peaks to an RD also allowed a quantitative estimate of their size by measuring the Feret diameter along the horizontal direction for each cluster of three or more peaks, which yielded a median size of 150 nm (Fig. 2G). We can therefore conclude that RDs have a typical diameter of 150 nm and typically contain four co-replicating stretches, which are separated by 63 nm, parameters well below the diffraction limit and consequently only detectable by super-resolution microscopy. Our findings provide a physical characterization of the internal structure of RDs *in situ* and are in line with previous biochemical estimates.

### Double labeling of neighboring RDs reveals their physical separation

Having characterized the internal organization of single RDs, we next addressed their spatial relationship within a chromosome. Previous reports based on diffraction limited microscopy estimated neighboring RDs to be very close or in direct contact with each other (Shaw et al., 2010), questioning the presence of linker domains. In order to address this question, we aimed to differentiate neighboring RDs by sequential labeling of the DNA backbone with two fluorophores. We introduced two pulses of labeled nucleotides into cells, the first one using ATTO-633-dUTP and the second one using ATTO565-dUTP (Fig. 1, 3). The pulses were separated by increasing time intervals (Δt = 0, 15, 30, 45, 60, 90 and 120 min; Fig. 3A), resulting in cells with double labeled CTs. We chose the time intervals taking into account previous reports of the progression of replication. Typically, RDs lying side-by-side in the genome are replicating consecutively during S-phase (Berezney et al., 2000; Cook, 1999). We assumed that genomically adjacent RDs were labelled by pulses 60 min apart, a well-documented replication timing in mammalian cells (Desprat et al., 2009). To measure the distance of neighboring RDs without perturbing chromatin, we acquired high-resolution 3D confocal images in living cells (Fig. 3B). As expected, the two signals co-localized if both nucleotides were introduced simultaneously, and became more separated with increasing time between pulses (Fig. 3B). We then determined the position of the diffraction limited signal peaks in 3D in both channels by computational image analysis and measured the nearest neighbor distances between the early and later labeled RDs. As expected, their distance increased with increasing pulse spacing, confirming the spatial progression of replication timing (Fig. 3C). For Δt=60 min, the reported time to complete replication of one RD, we measured a median RD distance of 300 nm (Fig. 3C). Considering our RD average diameter of 150 nm, this indicates that RDs may be physically separated by about 100 nm, meaning that co-replicating regions clustered in RDs are connected by intervening DNA sequences that do not contain any of those clusters and may thus be less compact.

**Fig. 3.**
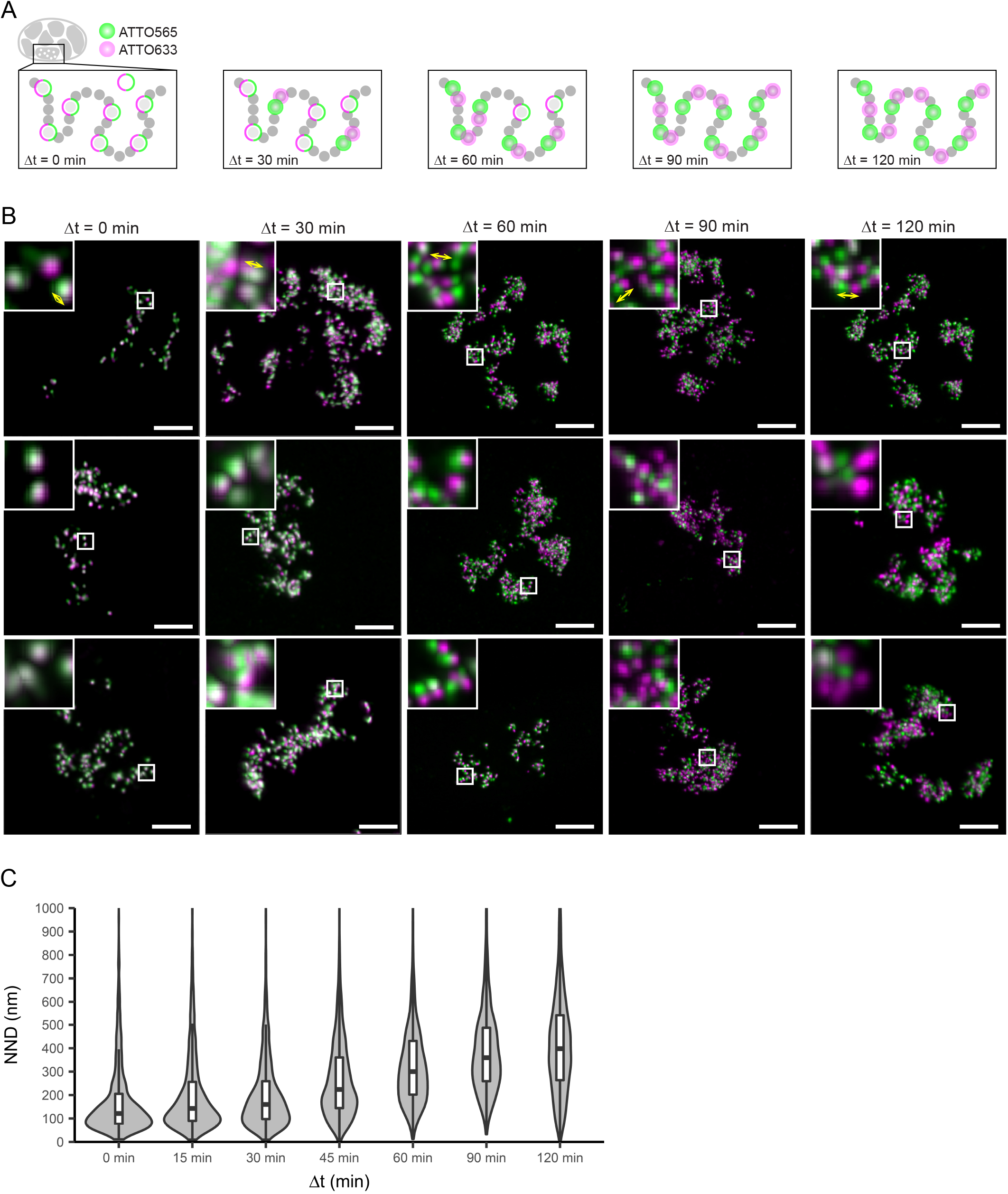
Dual-color confocal imaging shows neighboring domains spacing. (**A**) Schematics of the labeling pattern progression. A first round of co-replicative labeling with ATTO633-dUTP yields a first set of labeled RDs (green circles). A second labeling round with ATTO565-dUTP at increasing Δt targets either the same RDs or 1st / 2nd neighboring RDs, depending on Δt (magenta circles). The typical interval required to complete replication of one RD and proceed to the neighboring one is Dt = 60 min (Jackson and Pombo, 1998); therefore, we applied Dt = 0,15, 30, 45, 60, 90 and 120 min. (For the sake of space, Δt = 15 and 45 are not shown). (**B**) Images showing cells subjected to double-color labeling at Δt = 0, 30, 60, 90 and 120 min. Images correspond to maximum intensity projected z-stacks after deconvolution(50 nm pixels in xy, 150 nm pixels in z), and overlay of ATTO633 (green) and ATTO565 (magenta) channels. Scale bars 5 μm. Small boxes mark position of insets. Insets show 5x zoom-in detailed view of dual-color labeled RDs, with yellow arrows indicating the measured distances used to estimate median NND of neighboring RDs. (**C**) Violin plots showing the distribution and median NND between pairs of ATTO 633 and ATTO 565-dUTP labeled RDs at increasing Δt. Distances above 1000 nm are not shown but included in the determination of the quantiles. The median NND at Δt = 60 min was used as estimation on the nearest neighbor RDs spacing (*n*_NND_ = 2711 pair distances from 16 cells).

### Neighboring RDs are connected by flexible linkers

Being able to resolve the position of neighboring RDs in living cells, put us in a position to test how flexible or stiff they are connected along the chromatin fibre of a chromosome. To measure the mechanical coupling of neighboring RDs, we performed time-lapse microscopy of double-labeled CTs and analyzed their motion (Fig. 4, S2). If adjacent RDs were mechanically coupled, their movement should be correlated and the trajectories of neighbouring pairs be close to parallel. If adjacent RDs experienced little mechanical coupling due to less flexible linkers, their movement should be uncorrelated (Fig. 4A). We therefore assessed the degree of motion correlation by determining the mean correlation angle a between the displacement vectors of neighboring RD trajectories (with 0 degrees indicating complete coupling and 90 degrees no coupling; for details see Materials and methods).

**Fig. 4.**
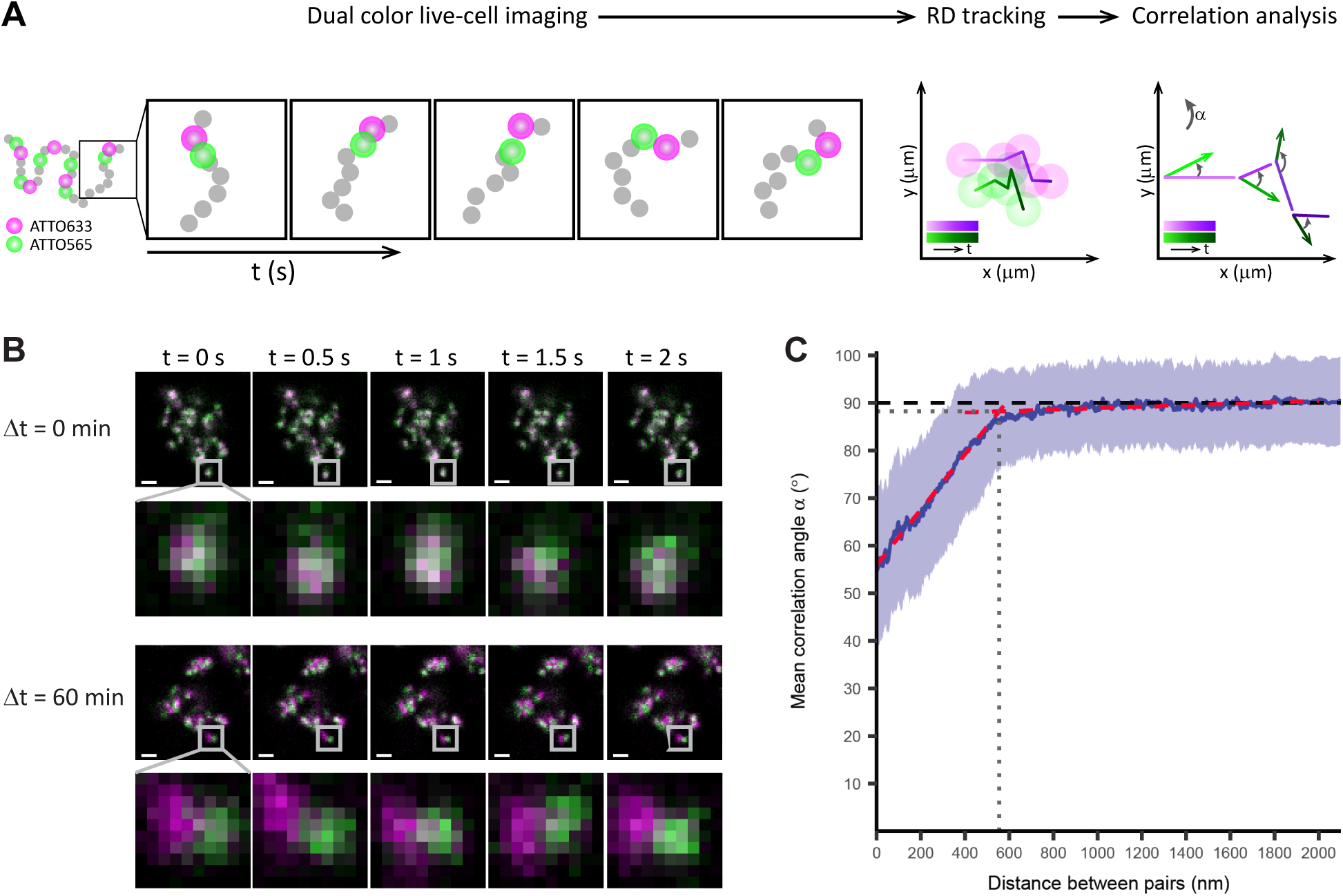
Dual-color live-cell confocal imaging detects a weak mechanical coupling of neighboring RDs. (**A**) Schematics of the experimental approach to measure the correlation angle a between trajectory pairs. A 2D temporal sequence (2 Hz, 30-300 frames) of dual-color labeled CTs is acquired (left, green: ATTO633-labeled RDs, magenta: ATTO565-labeled RDs). The position of each RD was determined and RD tracks were generated (middle). The correlation angle a is calculated at every time step from the corresponding displacement vectors (right). Then, for every trajectory pair, the mean a is calculated, and the averaged mean correlation angle <α> is extracted as a function of distance between pairs. To sample a broad distance interval trajectories from dual-color labeled RDs pulsed at increasing intervals between labeling pulses as described above, Δt = 0 min, 15 min (*n* = 3144 pairs, 26 cells), 30 min (*n* = 8256 pairs, 27 cells), 45 min (*n* = 5126 pairs, 21 cells), 60 min (*n* = 5306 pairs, 15 cells), 90 min (*n* = 4103 pairs, 22 cells), and 120 min (*n* = 4329 pairs, 22 cells) were acquired. (**B**) Time-lapse sequence of dual-color labeled RDs at Δt = 0 min (top), and 60 min (bottom). Scale bars 2 m. The 10x zoom-in panels show visually the correlated movement of RDs for Δt = 0 min and the uncorrelated movement of RDs for Δt = 60 min. (**C**) Density plot of averaged mean correlation angles for every trajectory pair <α> vs. the spatial distance (blue line <α>, SD shaded blue region). Bundled data from all Δt datasets (*n* = 30264 pairs, 133 cells). Two linear regression models were fitted respectively to data with distances < 400 nm and data with distances > 600 nm (red dashed lines). The transition point of correlated to uncorrelated movement was determined by the intersection of both (grey dotted lines).

High-resolution live-cell imaging and computational tracking of the motion of adjacent RDs sequentially labeled with two pulses of ATTO565- and ATTO633-dUTP spaced differently in time (Δt = 0, 15, 30, 45, 60, 90 and 120 min) covered a wide range of distances from below 100 nm to 1.5 μm (Fig. 4B, S2). Global motion correlation analysis of the data from a total of 36959 paired RD trajectories showed that moderate mechanical coupling was present at shorter pair distances while this coupling was rapidly lost with increasing pair distance (Fig. 4C). Fitting independent regression lines to the two linear coupling regimes apparent in the data identified a transition point at a distance of 550 nm and an angle of 88 degrees, close the 90 degrees expected for uncoupled motion (for details see Materials and methods). We can therefore conclude that adjacent RDs are loosely mechanically coupled at distances below 550 nm, which suggests that their linker sequences are highly flexible.

## Discussion

The structure and dynamics of chromatin in the mammalian cell nucleus remains poorly understood. Here, we addressed the internal organization, size, spacing and elastic connection of RDs using live and super-resolution microscopy. The key to achieve precise physical estimates of the structural features of RDs was the resolution provided by STORM and a fluorescent pulse labeling strategy that is restricted to co-replicating DNA stretches of RDs in single CTs after several rounds of mitosis. Our labeling approach (Fig. 1) causes minimal disruption of nuclear structure, the cells quickly recover from the scrape labeling procedure and it does not require harsh chemical treatments or DNA denaturation for introducing the fluorescent label.

With a median of four co-replicating stretches per RD, we found a slightly lower number than expected from previous reports, where the amount of replication forks per domain ranged from 6 to 20 (Berezney et al., 2000). Recently, single replicon imaging using structured illumination microscopy has been reported; however, this imaging approach has modest resolution improvement to approximately 100 nm and could therefore not resolve the fine details of the internal organization of RDs we measured here (Chagin et al., 2016). Our physical size estimate of 150 nm for RDs is expectedly significantly smaller than previous reports by confocal microscopy (Shaw et al., 2010), but in agreement with those reported by other super-resolution microscopies such as 3D-SIM (Baddeley et al., 2010), STED (Cseresnyes et al., 2009), and very recently by PALM (Nozaki et al., 2017). It is worth noting that our estimation likely represents a lower bound for the RD diameter, as the labeling is limited to short DNA stretches and the size is therefore based on the spatial arrangement of typically four active sites at the time of replication labeling and a total of approximately 70 kbp of labeled DNA. However, since the distribution of the active sites within the RDs at the time of labeling is likely to be random, and the labeled stretches maybe rearranged from their original position within the RD at the time of imaging, we deem our estimation a good representation of the general distribution of RD sizes.

Our improved resolution and double labeling approach also allowed us to estimate the physical distance of neighboring RDs to 300 nm, significantly larger than the median RD diameter of 150 nm. We recognize that the relationship of RD position and replication timing is still a matter of debate. Here, we assumed a timing gap of 60 min compatible with observations with population-based techniques (Desprat et al., 2009), and a locally sequential model of replication site progression consistent with single cell observations (Maya-Mendoza et al., 2010; Sporbert et al., 2002). Although the current literature is also consistent with a certain degree of stochastically initiated replication sites (e.g. Maya Mendoza et al., 2010), considering that we analyze a large number of physically neighboring RD pairs, our average conclusions should be a good representation of the behavior of RDs that lie next to each other on one chromosomal fiber. We thus propose that the average 150 nm gap between RDs is bridged by flexible DNA linker regions. Similar regions have been identified by HiC experiments as boundaries (Dixon et al., 2012) with genomic sizes of ca. 50 Kbp, compared to average RD/TAD size of 500 Kbp. Given that the physical size of RDs and linkers are similar, we can extrapolate that linker regions show an approximately tenfold lower degree of compaction compared to RDs and would therefore be expected to be much more flexible. Our analysis of mechanical coupling between neighboring domains in live cells confirmed this showing that coupling is largely lost at distances above 550 nm.

Based on the integration of the new structural and dynamic parameters of RDs determined in this study, we propose a new model for the *in situ* organization of chromosomal DNA (Fig. 5). Here, RDs constitute the stable structural units below the scale of the chromosome territory. They are on average 150 nm in diameter, spaced every 300 nm along the chromosome, and connected by flexible, less compact linkers of about 150 nm, which results in loose mechanical coupling of their motion that is largely lost beyond 550 nm. According to this model, flexible regions connect neighboring RDs that can thus move in a largely independent manner and will on average be spaced apart. Interestingly, the absence of mechanical coupling over longer length scales excludes the presence of stable “stiff” supra-structural units between a single RD and the scale of the chromosome territory, however we cannot exclude that transient highly flexible higher order arrangements may exist. Internally, each RD on average contains four co-replicating stretches, spatially separated by about 60 nm. The fact that the distance separating the co-replicating stretches remains constant across different sizes of RDs suggests a uniform packing of euchromatin RDs. This model provides an important basis for the quantitative study of chromatin organization in intact cells. Our labeling and imaging technology is sequence and cell-type independent and can be applied with minimal disruption of the native conformation. In the future, our approach can be used to address the reshaping of chromosomal structure and topology in processes such as cell differentiation or the formation of mitotic chromosomes.

**Fig. 5.**
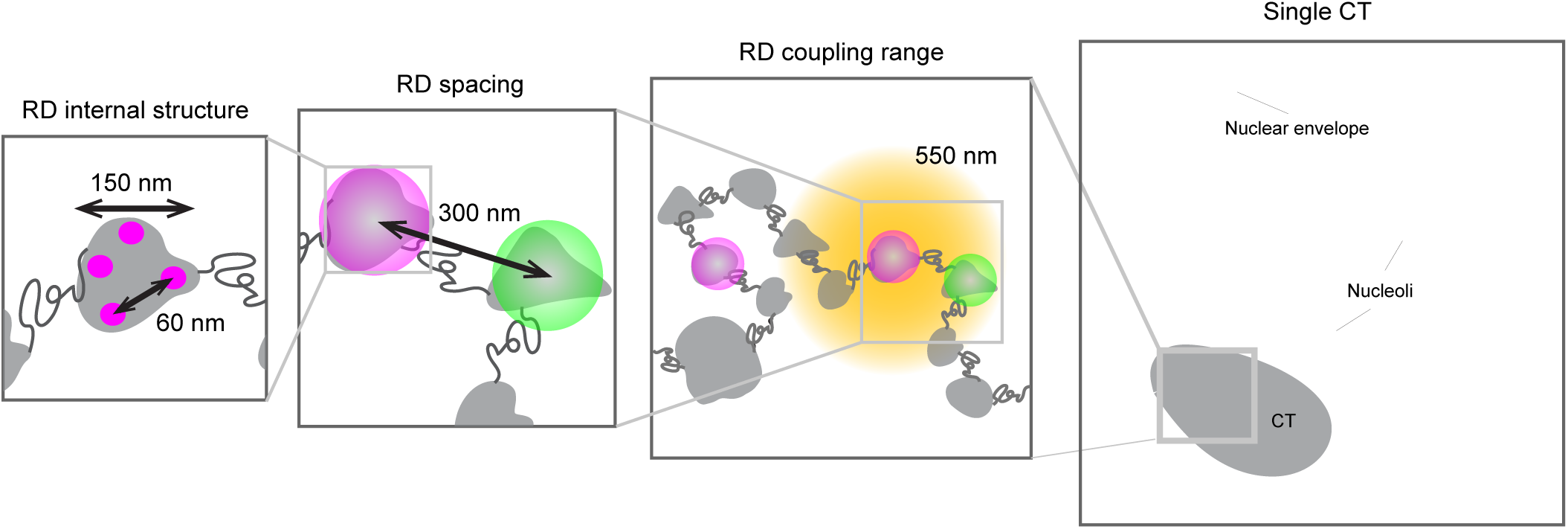
Summarizing model. The structural and dynamic information gathered from our experiments define a comprehensive model of higher order chromatin organization as follows: RDs median size is 150 nm and ranges up to approximately 400 nm. RDs comprise 4 co-replicating regions separated 60 nm on average. The typical nearest neighbor spacing (center to center) between RD is significantly bigger than typical RD size, with a median of 300 nm and ranges up to ca. 600 nm. We hypothesize the existence of extended linker regions between RDs with a median length of 150 nm. The elastic coupling range between RDs is lost at distances larger than 550 nm. RD = replication domain,= chromosome territory.

## Materials and methods

### Cell culture

We used normal rat kidney cells (NRKs) stably expressing PCNA-LAP BAC as pool (PCNA-LAP NRKs); the BAC construct was a kind gift from T. Hyman and I. Poser (MPI-CG, Dresden, Germany). The PCNA-LAP signal was not used in our study except in the initial setup of the replicative labeling protocol. PCNA-LAP exhibits a very specific change in nuclear distribution from G1 to S-Phase (from homogeneously distributed to punctate-like pattern) that can be monitored in confocal microscopy to optimize the time of application of the replicative labeling step after removal of the aphidicolin block (data not shown). Cells were cultured in high glucose Dulbecco’s modified Eagle’s medium (DMEM, Life Technologies) supplemented with 10 % v/v fetal bovine serum, 100 units/ml penicillin, 0.1 mg/ml streptomycin, 2mM glutamine, and 1 mM sodium pyruvate (complete medium). Cells were kept at 37 °C and 5% CO_2_.

### Co-replicative labeling of RDs

The labeling strategy consisted of three steps:

1. Cell synchronization by aphidicolin arrest at the G1/S transition: PCNA-LAP NRKs grown to 90% confluence were subjected to a mitotic shake-off. The suspension was collected, centrifuged at 200 x g for 4 min, and the supernatant was discarded. The harvested mitotic cells were plated in 8-well Labtek chambers (Thermofisher Scientific) with complete medium containing 1 μg/ml aphidicolin, and incubated for 10 h at 37 °C and 5% CO_2_.
2. Release of the aphidicolin arrest: Cells were washed 3x with 500 μl pre-warmed complete medium, and then incubated for 15 min to allow them to enter S-phase. A short waiting period of 15 min and a low aphidicolin concentration combined with the controlled arresting time, ensured proper recovery of the replication machinery (according to previous reports, 75% of forks are recovered after a 20 min waiting period [Davies et al., 2007]).
3. Pulse-labeling with a fluorescently labeled nucleotides: For single-color labeling, a staining solution of 15 μ of complete medium containing 67 μM ATTO 633-dUTP (ATTO-TEC GmbH) was added to S-Phase cells, then cells were scraped off the Labtek chamber with a rubber policeman, and incubated in the fluorophore solution for 1 min. Afterwards, fresh medium was added to a total volume of 500 μl. Cells with labeled single territories are obtained 5 days later, after several division rounds. For double-color labeling, the cells were exposed to 2 pulses of labeled nucleotides, namely ATTO 633-dUTP / ATTO 565-dUTP. The first pulse was introduced by scratching cells with a needle in the presence of ATTO 633-dUTP containing medium and incubated as described above. The second pulse with ATTO 565-dUTP was carried out at different times after the first pulse (Δt = 15, 30, 45, 60, 90 and 120 min), following the procedure described for single-color labeling. Although the scraping procedure detaches the cells from the chamber surface, it is very gentle, shown by high survival rates (>80%, from visual inspection of cells reattaching to the surface 6h after scraping). Survival rates after scratching for dual color labeling are ~70% prior to scraping (estimated by visual inspection of the scratched cells 6h after scratching).

### DNA combing and length calibration

We characterized our co-replicative labeling protocol by DNA combing (Parra and Windle, 1993; Michalet et al., 1997) on ATTO633-dUTP labeled cells. After the labelling procedure described above, cells were allowed to recover overnight in an 8-well Labtek chamber in complete medium at 37 C. Afterwards cells were trypsinized, harvested by centrifugation at ~400g for 5 min, and resuspended in PBS to a final density of 10^6^ cells/ml. 2 μl of the cell suspension was then spotted onto a silanized microscope slide, 10 mm below the slide frosted edge (Thermo Fisher), and lysed 10 min later with 5 μl of 0.5% SDS in 200 mM Tris-HCl, pH 7.4, 50 mM EDTA for 10 min at RT. After 15 min incubation at RT, the slide was placed vertically in a home built chamber containing 150 mM MES buffer pH 5.5 (DNA combing solution). The total volume of the chamber was 2 ml to ensure that the slide was immersed in the DNA combing solution, except for the frosted edge. The frosted edge was then attached to a home-built dip coater device (EMBL mechanical and electronic workshop), and the slide was pulled up at a constant speed of 300 μ.m/s. The dip coating procedure produces the linear stretching of the cells DNA onto the cover slide. After the slide was completely pulled up and out of the chamber, it was let to dry for 30 min at RT, and then fixed for 15 min in MetOH a-20 C. The sample was then washed 3 times with PBS, counterstained with PicoGreen following the protocol suggested by the manufacturer (Thermofisher), mounted in PBS and imaged by confocal microscopy to locate the ATTO633-dUTP signal coming from the replicative labeling pulse (Fig. S1C,E). A series of short (ca. 5-10 μ-m), comet-like stretches, were spotted along the PicoGreen stained fibrils. In order to convert the comet length into kbp, we calibrated the system by combing linearized, HindIII digested, λDNA fragments of 4.36, 9.41 and 23.13 kbp (NEB). We pre-labeled the fragments with PicoGreen and mixed them with a suspension of unlabeled, trypsinized NRK cells in the same concentration as utilized above. The λDNA-cell suspension was spotted as described before, and the slide was subjected to combing with the dip coater. After confocal imaging, the λDNA fragments were observed as straight, stretched segments along the glass surface. A linear fit of the data revealed a stretching rate of *s*_λDNA_ = 1.77 ± 0.5 Kb/μ.m (*n* >200 for each fragment length) (Fig. S1D, E), consistent with the value reported using different protocols (Shaw et al., 2010; Michalet et al., 1997). The resulting length of the ATTO633-labeled stretches was highly conserved with a mean of 18.2 ±4.5 Kb considering the λDNA calibration (Fig. S1F). We did a simple calculation to assess if the median number of 4 forks per RD is compatible with what is currently known about the process of replication. RDs typically finish replication within approximately 60 min (Jackson and Pombo, 1998; Ma et al., 1998). The replication speed of a single fork is in the order of 2 kb/min (Palumbo et al., 2013). Assuming 4 forks replicating DNA for 60 min, this would result in 480 kb of replicated DNA. This number matches the genomic length covered by one RD, which is approximately 500 kb according to the most recent systematic replication timing study on over 30 different cell lines (Pope et al., 2014). Formerly it was thought that RD are spanning bigger genomic regions, partially explaining the overestimation of typical numbers of forks per RD (Cook, 1999).

#### Confocal microscopy

Confocal imaging was carried out at a LSM 780 ConfoCor 3 microscope, equipped with a 63x 1.4 N.A. alpha Plan-Apochromat objective. GaAsP detectors were used for imaging PicoGreen in the DNA combing experiments. APD detectors were used for detecting signal from ATTO 633-dUTP and ATTO 565-dUTP.

#### DNA combing imaging

Imaging conditions for PicoGreen and ATTO633-dUTP were as follows: PicoGreen excitation at 488 nm, GaAsP emission collected between 490 and 550 nm; ATTO 633 excitation at 633 nm, emission long-pass filter 655 nm, and APD detection. Simultaneous scanning of both channels was carried out with a xy-pixel size of 90 nm; pixel dwell 8.16 μs / pixel, no averaging.

#### Live-cell imaging

Cells were plated onto 2-well Labtek chambers the day before imaging. Live-cell imaging was carried out at 37 °C in a CO_2_-independent medium (Life Technologies) supplemented with 20 % v/v fetal bovine serum, 100 units/ml penicillin, 0.1 mg/ml streptomycin, and 2 mM glutamine. To minimize photodamage, maximum light deposition of sample was kept below 5 J/cm^2^, and O_2_ concentration in the imaging medium was lowered to ~5% using the OxyFluor enzymatic system (Oxyrase Inc.). To this end, imaging medium was supplemented with Oxyrase (c_final_ = 0.3 U/ml) and sodium lactate (c_final_ = 10 mM, Sigma). O_2_ concentration was monitored with a FireSting O_2_ fiber-optic oxygen meter with a retractable needle tip (Pyro Science GmbH). Cells were incubated for at least 30 min before imaging to allow O_2_ to reach ~5%, and imaged up to 6 h.

Imaging conditions for the different fluorescent probes were as follows: LAP excitation at 488 nm, GaAsP emission collected between 490 and 550 nm; ATTO 633 excitation at 633 nm, emission long-pass filter 655 nm, and APD detection; ATTO 565 excitation at 561 nm, emission long-pass filter 545 nm, and APD detection; image voxel size x,y,z 90 x 90 x 400 nm, pixel dwell 8.2 μs, no averaging.

#### Timelapse 2D-confocal imaging for coupling range experiments

Simultaneous scanning of both channels was performed with the following parameters: xy-pixel size 90 nm; sampling rate 2 Hz; 30-100 frames were collected; image window 300×300 pixels; pixel dwell 8.16 μs per pixel, no averaging.

### Correlative confocal-STORM imaging

2D STORM experiments were carried out at a SR GSD microscope (Leica Microsystems) equipped with a HCX PLAPO 160 x, 1.43 N.A. Oil CORR-TIRF-PIFOC objective, a 642 nm laser for excitation (500 mW) and a 405 nm diode laser for back-pumping (30 mW). The lateral drift was minimized by the built-in Suppressed Motion (SuMo) stage. Images were acquired in epifluorescence mode with an Andor iXon3 emCCD camera (pixel size 100 nm, Andor) with 100 Hz acquisition rate. The microscope was equilibrated for 2 h before starting the experiments.

Cells were plated onto gridded coverslips (IBIDI) the night before imaging. The etched grid, lettered and numbered with 4 x 400 squares at 50 μm repeat distance, allows to image the cells first by confocal and then by STORM imaging. Confocal acquisition channel conditions for ATTO565- and ATTO633-dUTP were as described for live-cell imaging.

The day of the experiment, cells were fixed before imaging as follows: complete medium was removed, cells were washed twice with 1x PBS, and then fixed 4% PFA / 1x PBS (Electron Microscopy Science) for 30 min at room temperature. Finally, the PFA solution was removed by washing three times with 1x PBS.

After fixation and mounting in 1x PBS, confocal stacks were acquired on cells displaying 2-3 CTs (see confocal experimental conditions above). The imaged positions of the grid were annotated to allow an easy relocalization at the GSD microscope. The samples were then mounted for STORM in switching buffer (50 mM Tris-HCl pH 8.5 containing 10 mM NaCl, 10 mM MEA, 10 % w/v glucose, 0.5 mg/ml glucose oxidase, and 40 μg/ml catalase (all reagents from Sigma-Aldrich)) to ensure a consistent blinking of the fluorophores. STORM image acquisition was carried out as follows: the sample was irradiated with 642 nm laser at maximum power, to send most of fluorophores into a dark (non-emitting) state and achieve single molecule blinking (typically 20 s); after that, laser power was set to 60 % and a long series of images was acquired (30000-60000 frames), keeping the number of events detected rather constant (up to 15 events/frame) by manually increasing the 405 nm laser power to bring fluorophores back from the non-emitting state. Raw movies were saved for subsequent analysis.

A recent report illustrates the possibility of implementing STORM in 3D for replication foci and reliably address the drift over time, making it an useful tool for setups equipped with optical sectioning (Ma et al., 2017).

### Confocal analysis of RDs number and NND in dual color experiments

The position of RDs was determined from the 3D center of mass coordinates of molecules detected by the 3D ImageJ Particle Detector (Sbalzarini and Koumoutsakos, 2005). The center of mass measurements were performed in each fluorescence channel separately. Due to stochastic variation in two color labeling, this measure has a precision between the two channels for the same object around 120 nm, which is less than one RD diameter, but leads to non-zero distances for the case of simultaneous labeling of the same RD. This limited precision does not affect the average distance determination once the objects in the two channels are clearly separated after 30 min or longer labeling gaps, as the error is random for each object. The NND analysis between ATTO633- and ATTO565-labeled CTs was performed with a self-written Matlab 2012b (Mathworks Inc.) routine.

## Correlative confocal-STORM image analysis

### Preprocessing – Quality control

The localization of single molecule events was carried out with the Leica SR GSD Wizard. Briefly, a fast centroid fit was performed on pixels with intensity values above a set threshold (40 photons/pixel). The algorithm rejected peaks that were too bright, not circular, or too close to other peaks. Then, the integrated intensity for a single event was estimated from the number of photons (intensity/calibration factor of the camera, given by the manufacturer) collected after background subtraction. The wizard produces an event list with all localizations, which was exported in binary format and further processed in using custom-made routines in Matlab 2012b (Mathworks Inc.). The localization precision was 12 nm, estimated by statistical analysis of subdiffraction peak determination by Gaussian fitting. Three quality control steps were implemented to generate the final images: a photon count threshold to discard events with poor localization precision; a lateral drift correction to keep images with drift < 10 nm after correction; and finally, Fourier ring correlation to discard images with resolution worse than 40 nm. In detail, the quality control was performed as follows: The raw data was first visualized with the custom-made Matlab routine. Events with low localization precision (below 500 photons; conservatively determined from the photon count histogram) were filtered out. Afterwards, lateral drift was corrected by running a correlation-based routine (Szymborska et al., 2013). Finally, a super-resolution image was reconstructed from the events list by adding a single gray-value per localization event, with a pixel size of 10 nm based on the resolution of the microscope (20-30 nm). Since the STORM microscope was operated in epifluorescence mode, it did not provide optical sectioning and the excitation of fluorophores occurs effectively in an axial region ~1 μm thick across the focal plane. In order to identify individual CTs and select the corresponding regions in the image where fluorophores were effectively excited, the super-resolved image was overlayed with the corresponding confocal z-stack that contained the entire nuclear volume and labeled CTs. To perform the overlay, the confocal z-slices (x,y pixel size: 90 nm, z-interval 200 nm) were scaled up and flipped to match the dimensions and orientation of the STORM images (x,y pixel size: 10 nm). From the scaled images, a set of moving z-projections (sum intensity, sliding window = 5 slices) was generated throughout the stack. Each of the projected substacks was then registered against the STORM image using an iterative, intensity-based, rigid-body registration algorithm implemented in MATLAB 2012b. Afterwards, a normalized cross-correlation between each projected substack and the STORM image was computed, and the optimal projected substack was identified as the one with the highest score in the correlation matrix. From the resulting overlay, individual CTs in the STORM images were manually segmented (typically 2-3 CTs/nucleus), and images with individual CTs were generated. Regions in the STORM images coming from CTs outside the projected substack were cropped out to exclude out-of-focus structures with inherently poor resolution.

The experimental resolution was estimated by Fourier ring correlation analysis (FRC) (Nieuwenhuizen et al., 2013) on the individual CTs segmented from the overlay confocal-STORM, using the 3σ threshold criterion. Briefly, the event list for each CT was split into 2 temporal blocks, Fourier ring was calculated, and the radial profiles from the image center were plotted. When images had residual drift not detected as strong asymmetry in the Fourier-ring profile, such as rotational drift, this is reflected in the estimate for resolution. Images which showed an asymmetric Fourier-ring profile or displayed resolution worse than 50 nm were discarded. The images that passed the quality controls were then subjected to clustering analysis to quantitatively address the ultrastructure of the CRDs.

We also estimated the attainable resolution as traditionally informed, from the full-width at half maximum (FWHM) of the signal coming from isolated replicating DNA stretches in the super-resolved images (*n =* 25).

### Super-resolution analysis – clustering algorithm

The characterization of RDs was carried out on deconvolved images containing single CTs, applying a density-based clustering algorithm. First, a 3x3 median filter was applied and all pixels not connected to at least 3 other pixels were discarded. Then, individual RFs were detected applying grayscale dilation (radius 4 pixels). The identified replicons were clustered using the DBSCAN algorithm (Ester et al., 1996) using the following parameters MinPts = 2 and Eps = 14. From the replicon cluster analysis, the following parameters were calculated to characterize the structure of CRDs: 1) Total number of replicons, number of replicon clusters, number of solitary replicons; 2) Nearest neighbor (Euclidian) distance (NND) between replicons within clusters; and 3) Feret-Diameter of each cluster as a measure for RD size. The Feret-Diameter was selected to estimate the size of the RDs as they display irregular shapes (Feret, 1931). The Feret-Diameter gives the size of an object along a specified direction (here horizontal), and it can be defined as the distance between the two parallel tangential lines restricting the object perpendicular to that direction.

The STORM images also revealed the presence of unclustered peaks that were assigned to solitary replication stretches, not detectable in confocal microscopy due to their relatively low fluorescence signal. Although 2D-STORM was performed, the correlation with the confocal stack allowed us to exclude that these stretches are part of clusters, which have only been partially imaged because of the lack of z-sectioning capability of our super-resolution microscope. We excluded these structures from the clustering analysis, and we suggest that these stretches may correspond with previously reported origin-free, unidirectional forks (Palumbo et al., 2013).

## RDs coupling range

2D confocal time-lapse movies of double-color labeled CTs were acquired to quantify the loss of motion-correlation between RDs spaced at increasing distances from one another, using the correlation angle α between displacement vectors of trajectories in every time-step as estimate. A dataset using 7 different waiting times between nucleotide pulses from 0 minutes and to 120 min were recorded on a Zeiss LSM 780 ConfoCor and simultaneously scanned with both a 633 nm and a 565 nm-laser to excite RDs labeled with ATTO 633 and ATTO 565, respectively; the simultaneous labeling of Atto633-dUTP and Atto565-dUTP (Δt = 0 min) was used to assess the noise for the case of fully correlated RD movement; time-resolution was 0.5 s. Only euchromatic RDs were used for this analysis, since most heterochromatic RDs are immobilized. For data analysis, pairs of trajectories with track lengths of at least 15 s and up to 50 s were selected, the position of RDs was estimated using ImageJ, and <α> was then calculated using a self-written routine in Matlab. The dual-color labeling allowed us to determine the positions of RD pairs even if their signal overlapped, and enabled us to unambiguously track their motion with a precision of 30 nm. Two linear regression models were fitted respectively to data with distances < 400 nm and data with distances > 600 nm using the lm() function in R (R Core Team, 2017). The transition point was set to the intersection of the two regression lines.

We observed elastic coupling of RD rapidly decreasing with growing pair distance and largely lost at distance above 550 nm. This distance is significantly shorter than previously reported using image correlation analysis with fluorescently labeled histones where a temporal resolution of 10 s was used (Zidovska et al., 2013). To make our better time-resolved (0.5 s) data comparable, we re-analyzed it using a 10 s time step, but the 550 nm threshold at which correlation was lost remained unchanged. The difference in the observed values could be technical, due to the different methods and resolutions used, or biological, as histones will label both RDs and linker domains, while we specifically labelled only RDs.

## Online Supplementary Material

Figure S1 shows the characterization of co-replicative labeling of early-replicating RDs for correlative confocal-STORM imaging. Figure S2 shows individual time-points of the dual-color live-cell confocal imaging, which measures a weak mechanical coupling of neighboring RDs. Table S1 summarizes fluorophores tested for optimized replicative labeling of RDs using aminoallyl-dUTP derivatives in correlative confocal super-resolution imaging applications.

## Acknowledgements

We thank the EMBL Advanced Light Microscopy Facility (especially M Lampe and S Terjung) and EMBL mechanical and electronic workshop; A Szymborska and M Eltsov for advice with STORM and cell fixation; and the members of the Ellenberg group for fruitful discussions. The work was supported by the 4D Nucleome / 4DN NIH Common Fund (U01 EB021223) to JE and the European Molecular Biology Laboratory (WX, MJR, JKH, SA, JE). W.X. was supported by the EMBL International PhD Programme (EIPP). M.J.R. was supported by a Humboldt Foundation postdoctoral fellowship. S.H. was supported by an EMBO long-term Postdoc fellowship.

The authors declare no competing financial interests.

## Author contributions

W.X., J.R. and J.E. conceived the project and designed the experiments. W.X. and J.R. performed the experiments. S.H. established labelling assays. J.K.H. supported data and statistical analysis. J.R., W.X., S.A. and J.E. wrote the manuscript.

